# Machine learning-based revelation of GABBR1 and IQGAP2 as candidate biomarkers of pulmonary arterial hypertension and their correlation with immune infiltration

**DOI:** 10.1101/2023.10.27.564313

**Authors:** Bao Wang, Ju Cheng, Zengyou Li, Yanfeng Peng

## Abstract

Pulmonary arterial hypertension (PAH) is a pulmonary vascular disease with complex pathogenesis, and its intrinsic molecular mechanisms remain unclear. The aim of this study was to screen gene expression data from PAH patients, identify possible diagnostic indicators of PAH and to investigate the role of immune cell infiltration in the progression of PAH.

This study made use of the gene expression dataset of PAH patients from the GEO database. R software was used to identify differentially expressed genes and perform functional enrichment analysis. The SVM-RFE, LASSO and Random Forest algorithms were then used to screen for PAH hub genes and validated in the peripheral blood and lung tissue datasets. Finally, the CIBERSORT algorithm was used to assess PAH lung tissue immune cell infiltration and to investigate the correlation between hub genes and immune cells.

A total of 132 DEGs were screened in this study, which were centrally involved in the neuroreceptor-ligand activity pathway and associated with neurotransmission and hemoglobin complex. A total of 2 pivotal genes, GABBR1 and IQGAP2, were obtained by machine learning algorithms. The 2 pivotal genes had good predictive power as verified by ROC curves. Further immune infiltration analysis showed a decrease in T cells and an increase in the proportion of macrophages and dendritic cells in the lung tissue of PAH patients. The expression of GABBR1 was positively correlated with T cells and negatively correlated with macrophages and dendritic cells.

In our study, we identified 2 potential diagnostic key genes: GABBR1 and IQGAP2. Our findings may provide a theoretical basis for the analysis of the underlying mechanisms of PAH and the development of targeted medicines.

**Highlight Box:** *Key findings:*

- We identified 2 potential key genes of PAH, GABBR1 and IQGAP2.

*What is known and what is new?:*

- Sympathetic hyperexcitability as well as immune responses are closely associated with the development of PAH, and pulmonary vascular hyperplasia is a key pathogenetic mechanism of PAH.
- Important biomarkers related to neuroreceptors and immune responses in PAH lung tissue have not been identified, while our study identified GABBR1 as a key neuroreceptor and immune cell regulator in PAH. IQGAP2 could be a new hotspot direction for pulmonary vascular remodeling.

*What is the implication, and what should change now?:*

- GABBR1 and IQGAP2 may be potential therapeutic targets for PAH. The new horizon provided by this study will provide some reference for subsequent PAH studies.

## 1. Introduction

Pulmonary arterial hypertension (PAH) is a complex pulmonary vascular disease characterized clinically by elevated pulmonary artery pressures and downstream hemodynamic abnormalities in the pulmonary vasculature and right ventricle (1). Due to this characteristic, resting mean pulmonary artery pressure (mPAP) ≥20 mm Hg had been mostly used as an important diagnostic criterion in recent studies (2). The current global prevalence of PAH is approximately 1% (3). The symptoms of PAH onset are non-specific and patients usually present with dyspnea on exertion (4). However, PAH is a life-threatening disease and without effective intervention, pulmonary hypertension usually progresses to right ventricular failure and death (5). Pulmonary viral infections are strongly associated with the development of PAH, and available studies have confirmed that infection with coronavirus (COVID-19) can lead to pulmonary vasculopathy, suggesting an increased chance of PAH in the future (6). However, the specific mechanisms underlying the pathogenesis of PAH are complex, and the mechanisms underlying the transformation of normal lung tissue to lesions remain unclear. Therefore, understanding the core biomarkers of PAH is essential for the diagnosis and treatment of this complex disease.

PAH has a high correlation with the patient’s living environment, metabolism, genetics and hypoxic injury, with the activation or inhibition of molecular mechanisms resulting from many factors (7). Among these are dysregulation of pathways such as oxidative stress, inflammation and metabolic dysregulation, which ultimately lead to pathological changes such as vasoconstriction, smooth muscle dysregulation and endothelial cell proliferation in the pulmonary vasculature (8). It has been shown that BMPR2, a key gene for endothelial cell proliferation, is the most common gene mutated in hereditary PAH, and by upregulating its expression, the PAH process can be effectively interfered with (9). However, mutations in BMPR2 are less likely to occur in idiopathic PAH (10). The above findings suggest that exploring the core regulatory genes may be a novel therapeutic strategy for PAH.

Immune cells and inflammatory factors are among the important factors contributing to the altered pulmonary vascular pathology in PAH (11). In patients with PAH there is a significant decrease in T-cell function, along with altered composition ratios of monocytes, macrophages, dendritic cells, natural killer cells and B cells (12). In thymus-free nude mice congenitally lacking T cells, their lungs are infiltrated with macrophages, mast cells and B cells, exhibiting immune infiltrative features similar to those of PAH lesions (13). Inflammatory factors, such as IL-6, have an important role in PAH development. mice with IL-6 overexpression spontaneously develop PAH and pulmonary vascular remodeling, as well as occlusive remodeling in hypoxia similar to human disease, whereas IL-6 knockout mice are more resistant to the development of PAH induced by chronic hypoxia (14). The above findings suggest that studies targeting immune-related genes may provide new perspectives for the treatment of PAH.

In this study, we obtained biochip data from the Gene Expression Omnibus (GEO) database to analyze differentially expressed genes (DEGs) (15). Machine learning algorithms were used to further screen and identify diagnostic factors for the PAH hub gene and to validate the diagnostic performance of the core gene by ROC analysis. Further we used CIBERSORT algorithm to study differential immune infiltration between 22 immune cell subpopulations in lung tissue of PAH patients (16). In addition, to better understand the potential molecular immune mechanisms of hub genes, we investigated the relationship between hub genes and infiltrating immune cells. We present the following article in accordance with the TRIPOD reporting checklist.

## 2. Methods

### Data download

The GSE15197, GSE33463 and GSE113439 datasets (GPL6480 platform, GPL6947 platform and GPL6244 platform) were downloaded from the GEO database. In each dataset, only samples from normal control patients (Con group) and PAH patients (PAH group) were selected. GSE15197 contains 31 lung tissue samples, 13 from normal control patients and 18 from PAH patients. GSE113439 contains 26 lung tissue samples, 11 from normal control patients and 15 from PAH patients. GSE33463 contains 71 peripheral blood samples, 41 from normal control patients and 30 from PAH patients. The study was conducted in accordance with the Declaration of Helsinki (as revised in 2013).

### Data processing and consolidation

The GSE15197, GSE33463 and GSE113439 datasets were annotated using R 4.3.0 software (https://www.r-project.org/) and duplicate values were removed using the mean method. The “inSilicoMerging” package was used to merge the GSE15197 and GSE113439 datasets, and the “limma” package was further used to remove batch effects. The combined dataset consisted of 57 samples, 24 from normal control patients and 33 from PAH patients, with 16,869 genes tested. This dataset will be used for subsequent studies. the GSE33463 dataset will be used for subsequent validation.

### DEGs identification

Use the “limma” package of R 4.3.0 software to filter and identify differentially expressed genes (DEGs) in the combined dataset. A threshold of |LogFC| > 1 and Adjusted *P* value < 0.5 was set. The results obtained were visualized using the “ggplot2” package. On this basis, the “pheatmap” package was used to draw the heat map, setting the clustering method to “complete” and the distance algorithm to “eulidean”. To highlight the differences between the Con and PAH group, we used the “vegan” package to plot PCoA and set the difference algorithm to “anosim”. DEGs were used for follow-up studies and undefined genes were removed.

### Functional enrichment analysis

We used the DIVID database (https://david.ncifcrf.gov/) to obtain the biological functions and signaling pathways involved in the DEGs. *P* value < 0.05 was set to indicate significant functional enrichment. Revealing the potential biological functions of DEGs by GO analysis. Potential molecular pathways used to analyze core gene action by KEGG analysis. Demonstrate the association of DEGs with disease by Disgenet analysis. All results were visualized using the “ggplot2” package of the R 4.3.0 software.

### Hub gene screening by machine learning

We used a combination of SVM-RFE, LASSO and Random Forest algorithms for the screening and identification of pivotal genes. The SVM-RFE algorithm was implemented using the “e1071” and “caret” packages of the R 4.3.0 software. The cross-validation method was set as k-fold cross-validation, divided into 10 folds and repeated 5 times. The LASSO algorithm was implemented using the “glmnet” package of the R 4.3.0 software, using “lambda_min” to filter the best set of variables. The Random Forest algorithm was implemented using the “randomForest” package to build 10,000 decision trees to obtain the pivot genes, and the results were considered in terms of both the increase in MSE (%incMSE) and increase in NodePurity (incNodePurity) metrics. The results of the three machine learning algorithms use the “venneular” package to take the intersection and obtain the hub genes.

### Hub genetic diagnostic performance validation

We validated the pivotal gene diagnostic performance using the “pROC” package of R 4.3.0 software. Further logistic regression was performed using the “glm” function, line-column plotting was performed using the “nomogramFormula” package, the accuracy of the model was assessed based on the consistency index, and calibration curves were plotted to assess the performance of the model.

### Analysis of immune cell infiltration

Immune cell infiltration studies were performed using the “CIBERSORT” package in R 4.3.0, with the objective of obtaining a matrix of immune cell infiltration in lung tissue from PAH patients. The immune cell infiltration matrix data was visualized using the “ggplot2” package and heat maps were drawn using the “pheatmap” package to highlight the differences in immune cell infiltration between the Con and PAH group. The wilcoxon rank sum test was used to analyze the differences between the two groups in terms of immune cells.

### Analysis of the correlation between hub genes and immune cells

Correlation of hub genes with infiltrating immune cells was calculated using Spearman’s correlation coefficient. The “ggpubr” package in R 4.3.0 was applied for visualization.

### Prediction of regulatory hub gene miRNAs

Regulatory microRNAs (miRNAs) for hub genes were obtained from the mirnet database (https://www.mirnet.ca/). The search results were visualized by using the “ggalluvial” package of the R4.3.0 software.

### Statistical analysis

All statistical analysis for this study were implemented using R 4.3.0 software, and all statistical tests were bilateral. Statistical analysis of the two data sets was performed using the wilcoxon rank sum test. Graphical visualization was implemented using the “ggplot2, ggpubr and pheatmap” packages. *P* < 0.05 was considered statistically significant.

## 3. Results

### Data pre-processing and DEGs identification

Two datasets, GSE15197 and GSE113439, were included in this study and the two datasets were combined and corrected (*Figure* 1A, B). PCoA analysis of the Con and PAH groups in the combined dataset revealed significant differences between the two (*Figure* 1C), suggesting that the correction operations performed on the dataset did not affect their original differences. The results of the volcano plot showed that a total of 132 DEGs were obtained in the combined dataset, including 103 up-regulated and 29 down-regulated genes (*Figure* 2A). The clustering heat map showed that there were significant differences in expression between the Con and PAH groups on these DEGs (*Figure* 2B).

**Figure 1.**
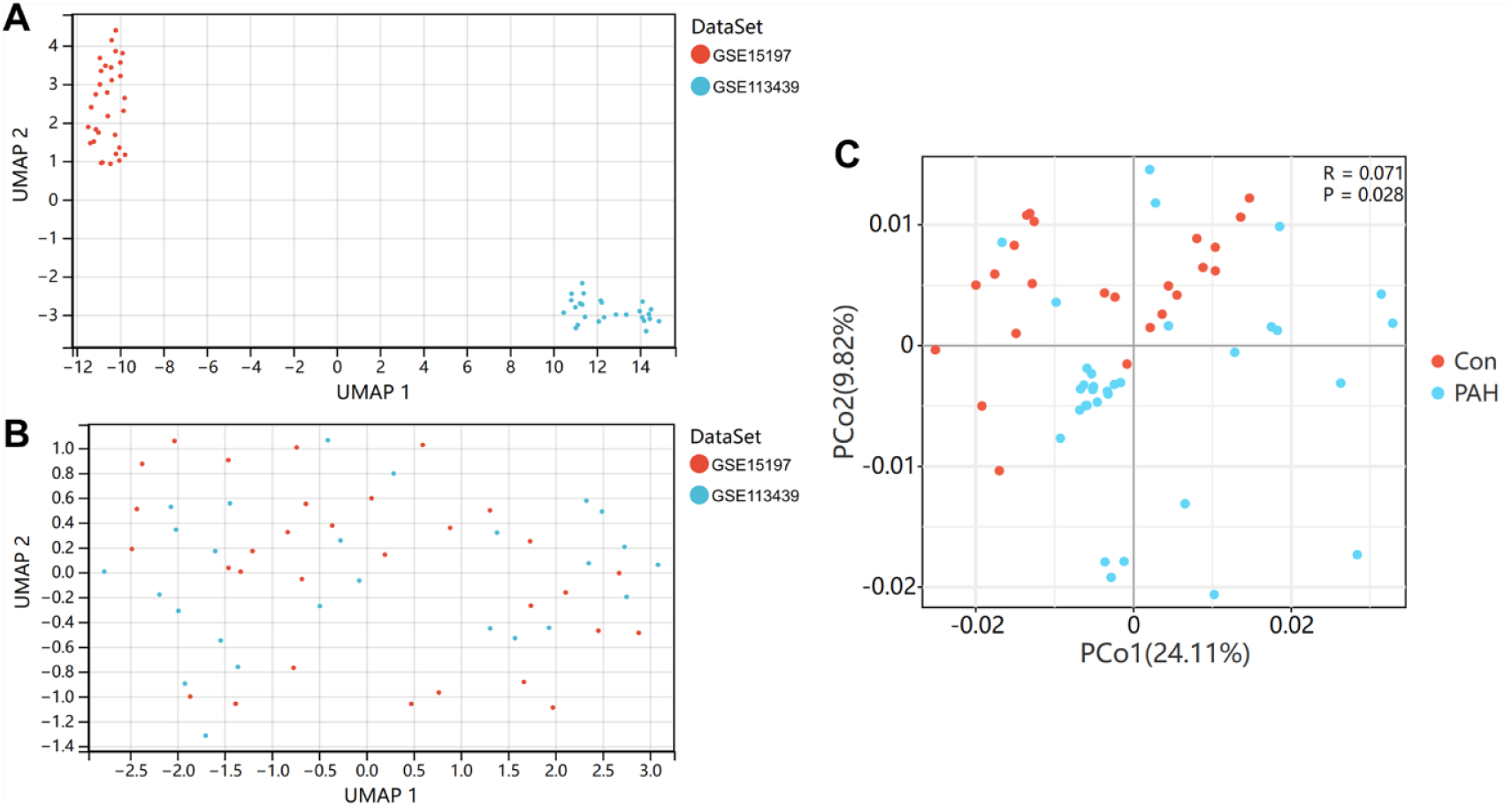
Normalization of merged data sets. (A) UMAP plot before data set normalization; (B) UMAP plot after data set normalization; (C) PCoA plot of the Con group and PAH group. PAH, pulmonary arterial hypertension.

**Figure 2.**
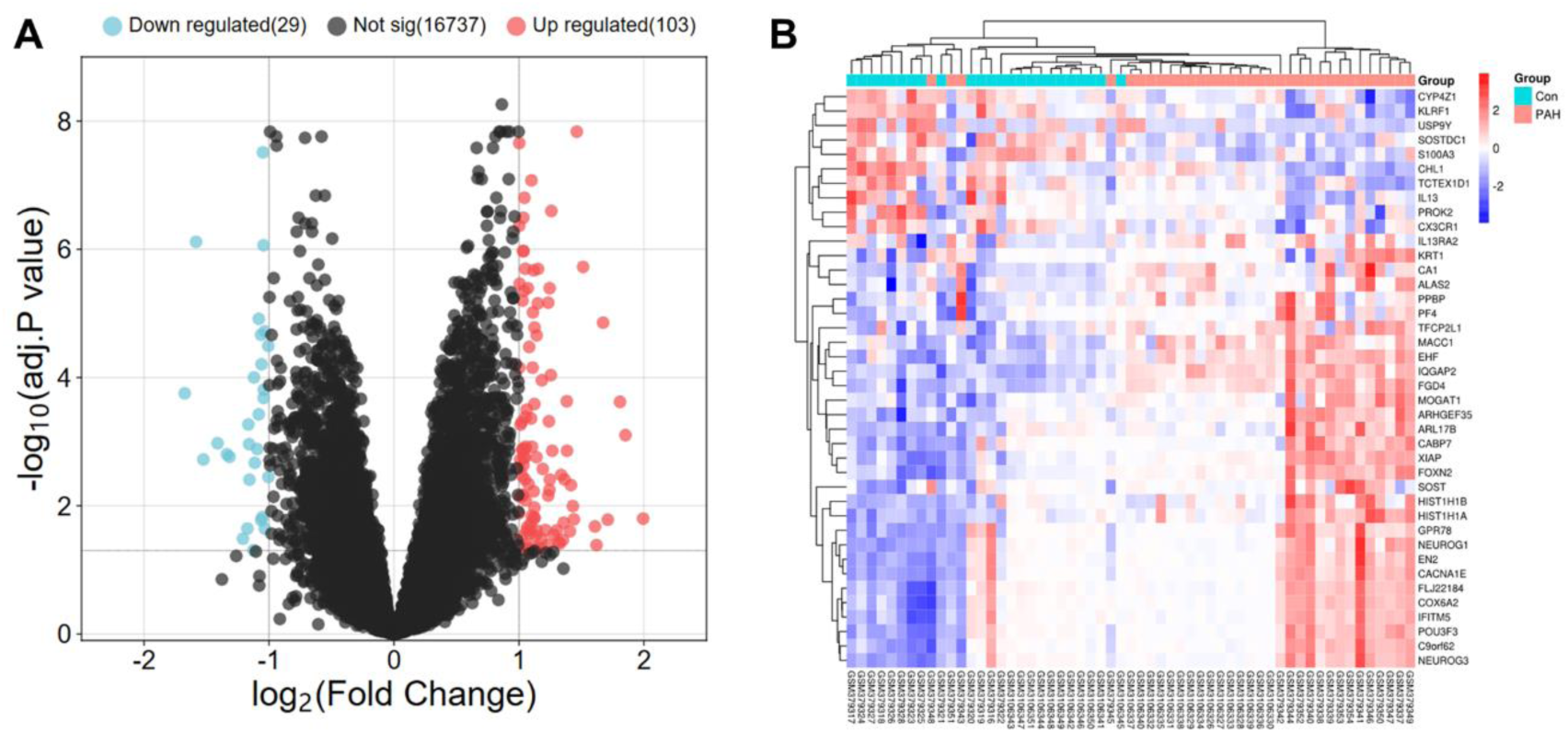
Volcano map and heat map of DEGs expression of PAH. (A) Volcano plot of DEGs (The differences are set as |LogFC| > 1 and Adjusted *P* value < 0.5); (B) Heatmap of DEGs expression (Top 40 up-regulated and 10 down-regulated genes). DEGs, differentially expressed genes; PAH, pulmonary arterial hypertension; FC, fold change.

### DEGs functional enrichment analysis

We performed KEGG analysis on the DEGs and found that the DEGs were mainly involved in four pathways: Taste transduction, Viral protein interaction with cytokine and cytokine receptor, Cytokine-cytokine receptor interaction and Neuroactive ligand-receptor interaction (*Figure* 3A). Further analysis revealed that GABBR1 in DEGs is mainly involved in Taste transduction and Neuroactive ligand-receptor interaction pathways. Its significant downregulation in the PAH group suggests that the physiological activity of neural ligand and receptor molecules in lung tissue of PAH patients may be attenuated. GO analysis showed that DEGs are associated with the regulation of neurogenesis, oxygen transport and neuropeptide signaling pathway in biological processes. In terms of cellular composition, they are highly associated with the hemoglobin complex, haptoglobin-hemoglobin complex and extracellular space. Molecularly involved in organic acid binding, haptoglobin binding and RNA polymerase II transcription factor activity and sequence-specific DNA binding. These results suggest that DEGs are primarily involved in the neural activity of lung tissue and are highly correlated with oxygen transport (*Figure* 3B). We further analyzed the association of DEGs with diseases through the Disgenet database. It was found that DEGs were highly associated with lung diseases such as Pneumonitis and Lobar Pneumonia, but also with immune responses such as Allergic Reaction, suggesting that immune responses have a large impact in PAH. The significant association of DGEs with diseases such as cardiac arrhythmias explain the molecular basis for the subsequent migration of PAH to cardiac diseases such as right ventricular failure (*Figure* 3C).

**Figure 3.**
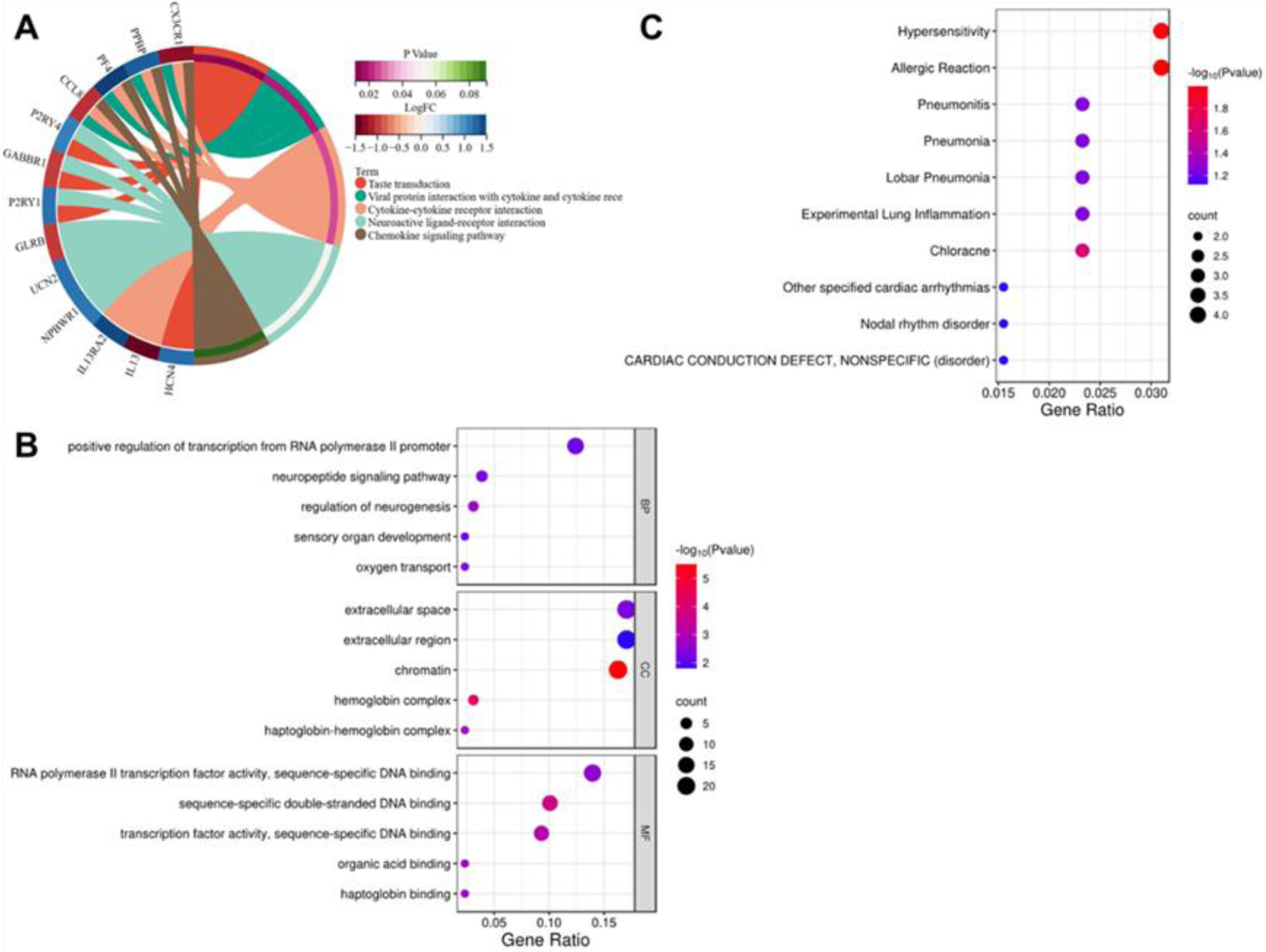
KEGG, GO and Disgenet analysis of DEGs. (A) KEGG analysis. (B) GO analysis (Top 5 according to *P* value in BP, CC, and MF, respectively). (C) Disgenet analysis (Top 10 according to *P* value in Disgenet). DEGs, differentially expressed genes; GO, Gene Ontology; KEGG, Kyoto Encyclopedia of Genes and Genomes; BP, biological processes; CC, cellular components; MF, molecular functions.

### Identification of hub genes

We used the SVM-RFE algorithm for optimal gene subset screening and found that the total dataset of 132 DEGs had the lowest RMSE (Figure 4A). On this result, we called up the top 10 genes in importance as pivotal genes (Figure 4B). There are 15 hub genes in the “lambda_min” subset obtained by the LASSO algorithm (Figure 4C, D). The Random Forest algorithm took the top 10 genes in the %incMSE and incNodePurity items as the optimal subset, respectively (Figure 4E, F). The intersection of these two optimal subsets was taken and a total of 9 hub genes were obtained. The three machine learning results were combined and a total of 2 hub genes were identified: GABBR1 and IQGAP2 (*Figure* 4H). Validation in the combined dataset revealed that GABBR1 expression was significantly higher in the Con group than in the PAH group (*P*< 0.001), while IQGAP2 showed a trend towards significantly lower levels (*P*< 0.001) (Figure 4I, J).

**Figure 4.**
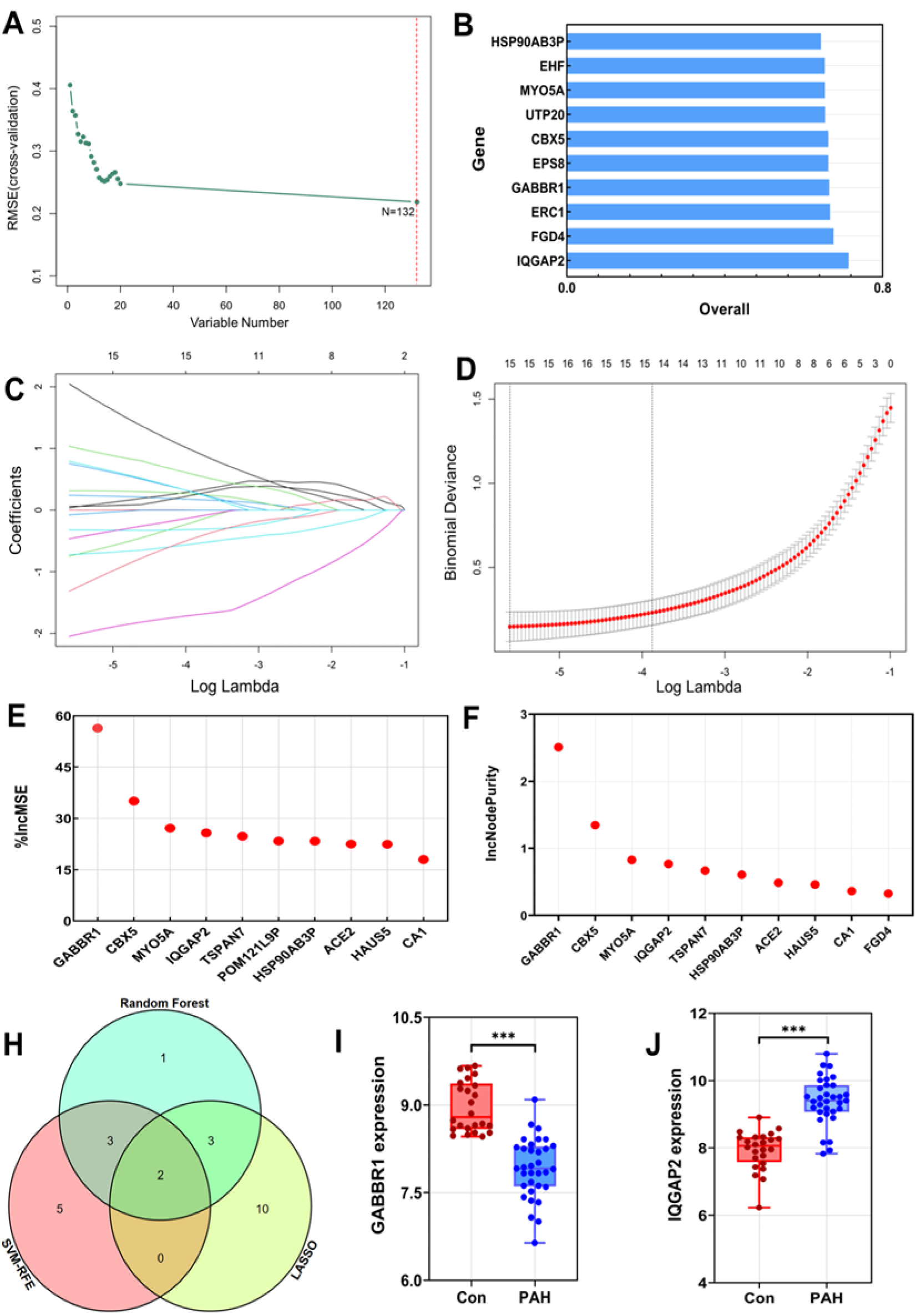
SVM-RFE, LASSO and Random Forest algorithms for screening hub genes. (A) RMSE values for different subsets of the number of variables in the SVM-RFE algorithm. (B) Top 10 variables in the optimal subset of the SVM-RFE algorithm. (C) Coefficient distribution of genes in the LASSO algorithm. (D) Distribution of Lambda values for different subsets of the LASSO algorithm. (E) Top 10 genes with %incMSE values in Random Forest algorithm. (F) Top 10 genes with IncNodePurity values in Random Forest algorithm. (H) Venn diagram of three machine learning algorithms. (I) Expression plot of GABBR1 in the combined dataset. (J) Expression plot of IQGAP2 in the combined dataset. SVM-RFE, recursive feature elimination for vector-holding machines; LASSO, least absolute shrinkage and selection operator; RMSE, root mean squared error; %incMSE, increase in MSE; IncNodePurity, increase in NodePurity; ***, *P*< 0.001.

### Hub genetic diagnostic efficacy validation

We validated the diagnostic efficacy of the pivotal genes obtained from machine learning screening by ROC curves. Firstly, the AUC values of GABBR1 and IQGAP2 were found to be 0.962 (0.915-1.000) and 0.933 (0.867-0.999) in lung tissue (combined gene set), respectively, and their combined validation resulted in an AUC value of 0.996 (0.988-1.000) (Figure 5A). Validation of the gene expression set in peripheral blood (GSE33463) revealed AUC values of 0.676 (0.542-0.809) for GABBR1 and 0.530 (0.390-0.670) for IQGAP2, respectively, and after combining them for validation, their AUC values were 0.661 (0.531-0.791) (Figure 5B). In the combined dataset, a corrected C index of 0.999 was constructed for the model. line column plots suggest that reduced GABBR1 expression and upregulated IQGAP2 expression have a greater risk of disease (Figure 5C). In the peripheral blood dataset, a corrected C index of 0.751 was constructed for the model. line plots indicate that IQGAP2 expression has greater diagnostic efficacy compared to IQGAP2 expression (Figure 5D). These results suggest that GABBR1 and IQGAP2 are diagnostic in both peripheral blood and lung tissue, and that the combination of the two genes provides better diagnostic efficacy.

**Figure 5.**
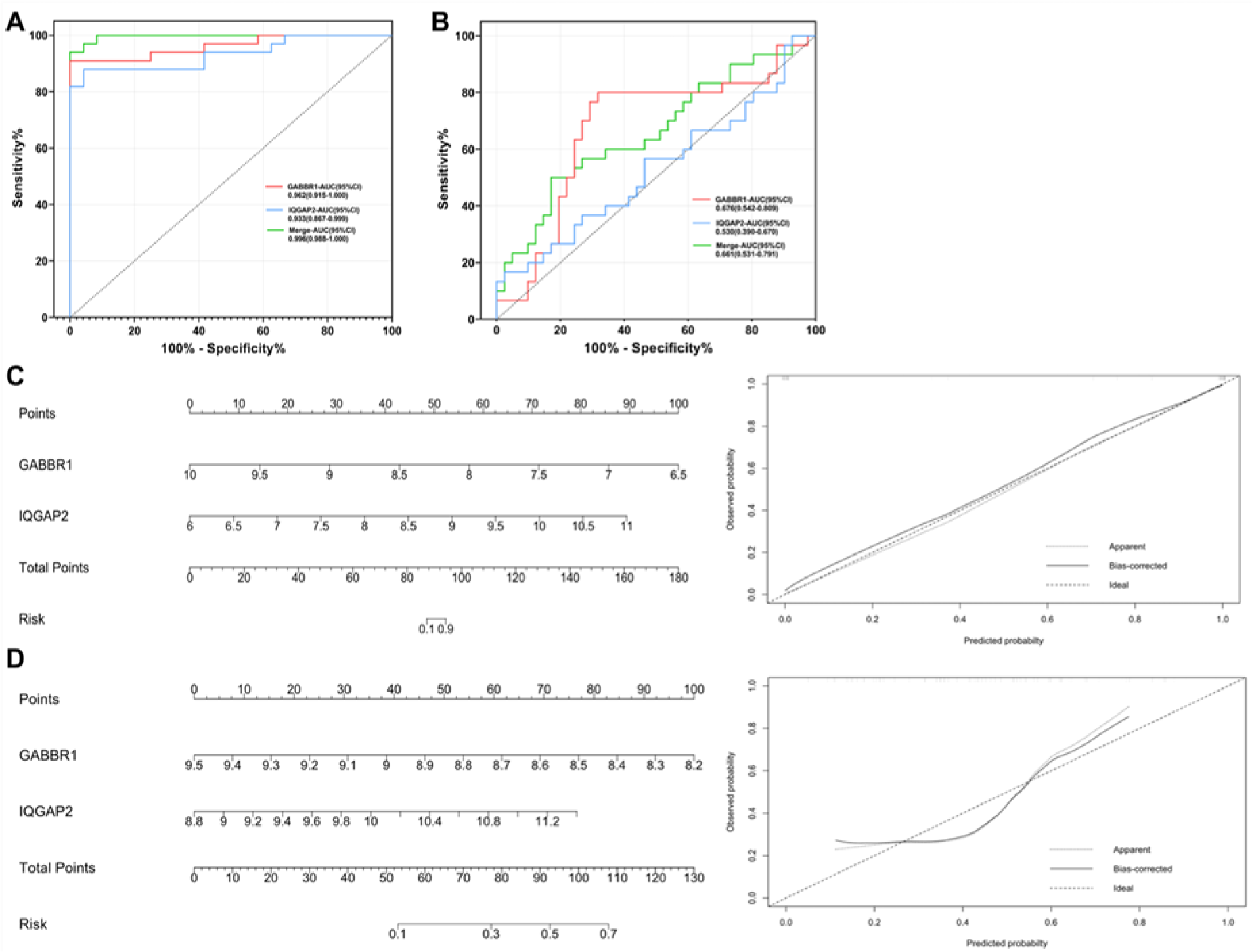
Validation of the diagnostic efficacy of GABBR1 and IQGAP2. (A) ROC curve of combined data set. (B) ROC curve of GSE33463 data set. (C) Nomogram column line chart of combined data set. (D) Nomogram column line chart of GSE33463 data set. ROC, receiver operating characteristic.

### Immuno-infiltration analysis

Differences in immune infiltration between normal and PAH lung tissue in 22 immune cell subpopulations were studied using the CIBERSORT algorithm. Immune cell stacking plots showed significant population differences between normal controls and PAH patients (Figure 6A). In normal lung tissue, T cells CD4 naive and T cells follicular helper had a predominant share. In PAH lung tissue, however, the proportion of Dendritic cells resting, Dendritic cells activated and Macrophages M0 was substantially increased. The clustered heat map showed that the normal group of lung tissue showed a positive correlation with immune cells of the T cell class, such as T cells follicular helper and T cells CD4 naive. In addition, the immune environment of normal lung tissue is mostly composed of Plasma cells and Neutrophils. PAH lung tissue, in contrast to the above trend, showed a substantial decrease in the T-cell content of its immune microenvironment and an increase in the content of dendritic cells and macrophages (Figure 6B). By comparing two with two, we get a clearer variation in the differences. The results showed that the levels of T cells CD4 naive (*P*= 0.001), T cells CD4 memory resting (*P*= 0.002), T cells CD4 memory activated (*P*= 0.002) and T cells follicular helper (*P*= 0.001) were significantly higher in the lung tissue of the normal group than in the PAH group. In contrast, the levels of Macrophages M0 (*P*= 0.032), Mast cells resting (*P*= 0.006) and Mast cells activated (*P*= 0.013) were significantly higher in the PAH group (Figure 6C).

**Figure 6.**
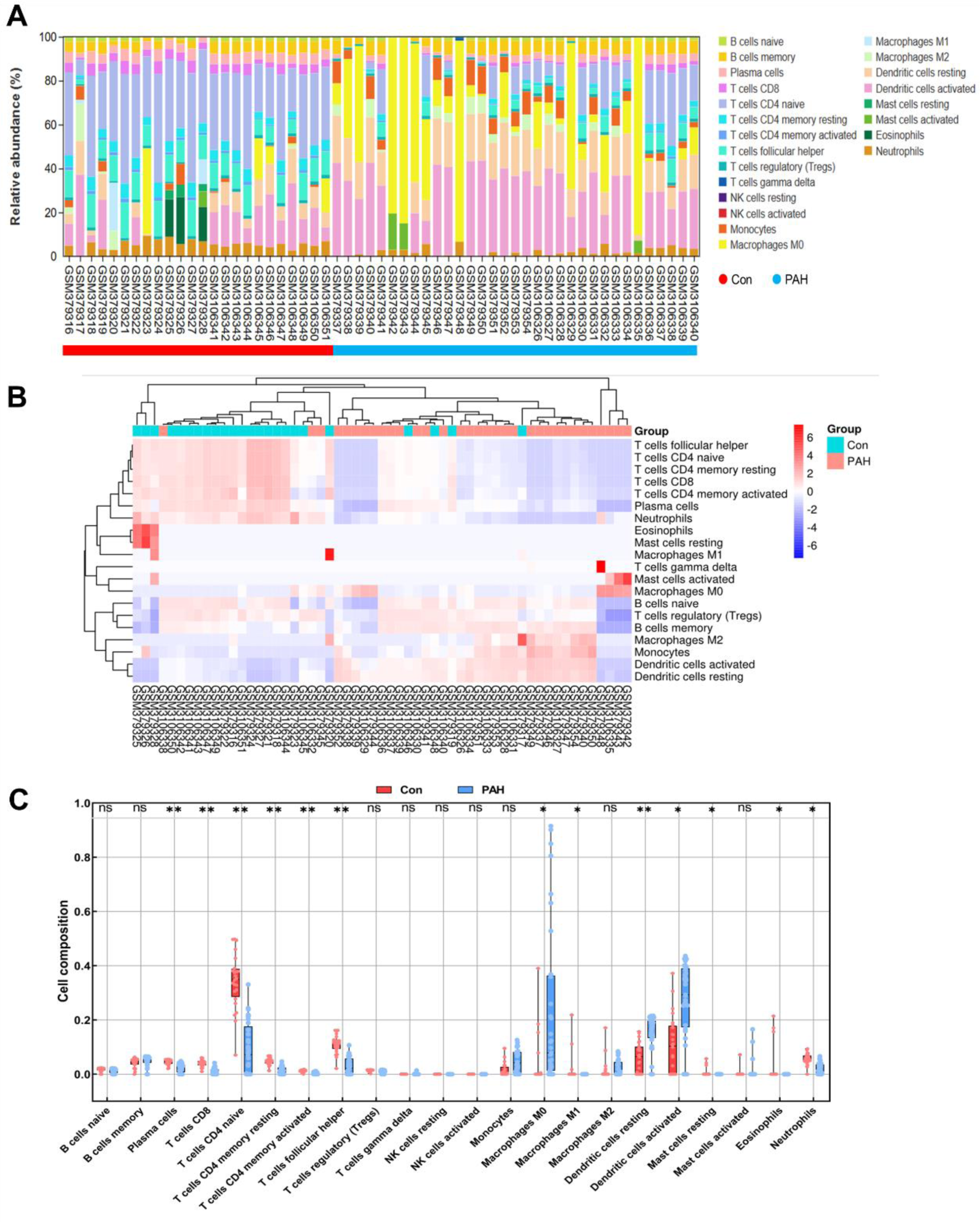
Immune infiltration analysis of the combine data set. (A) Immune cell accumulation plot. (B) Heat map for immune cells. (C) Box plot of immune cell infiltration levels in healthy individuals and PAH patients. PAH, pulmonary arterial hypertension; ns, *P*> 0.05; *, *P*< 0.05; **, *P*< 0.01.

### Analysis of the correlation between hub genes and infiltrating immune cells

Our results showed that T cells CD4 naive was significantly and positively correlated with T cells follicular helper (R= 0.997, *P*< 0.001), T cells CD4 memory resting (R= 0.997, *P*< 0.001), T cells CD4 memory activated (R= 0.925, *P*< 0.001) and Neutrophils (R= 0.804, *P*< 0.001) were significantly and positively correlated. There was a significant negative correlation with Macrophages M0 (R= −0.696, *P*< 0.001), Dendritic cells resting (R= −0.624, *P*< 0.001) and Dendritic cells activated (R= −0.557, *P*< 0.001). IQGAP2 was weakly associated with immune cells among the two pivotal genes. However, GABBR1 was significantly and negatively correlated with Macrophages M0 (R= −0.543, *P*< 0.001), Dendritic cells resting (R= −0.591, *P*< 0.001), Dendritic cells activated (R= −0.529, *P*< 0.001). Significant positive correlations were found with T cells CD4 naïve (R= 0.834, *P*< 0.001), T cells CD4 memory resting (R= 0.824, *P*< 0.001) and T cells follicular helper (R= 0.835, *P*< 0.001) (Figure 7).

**Figure 7.**
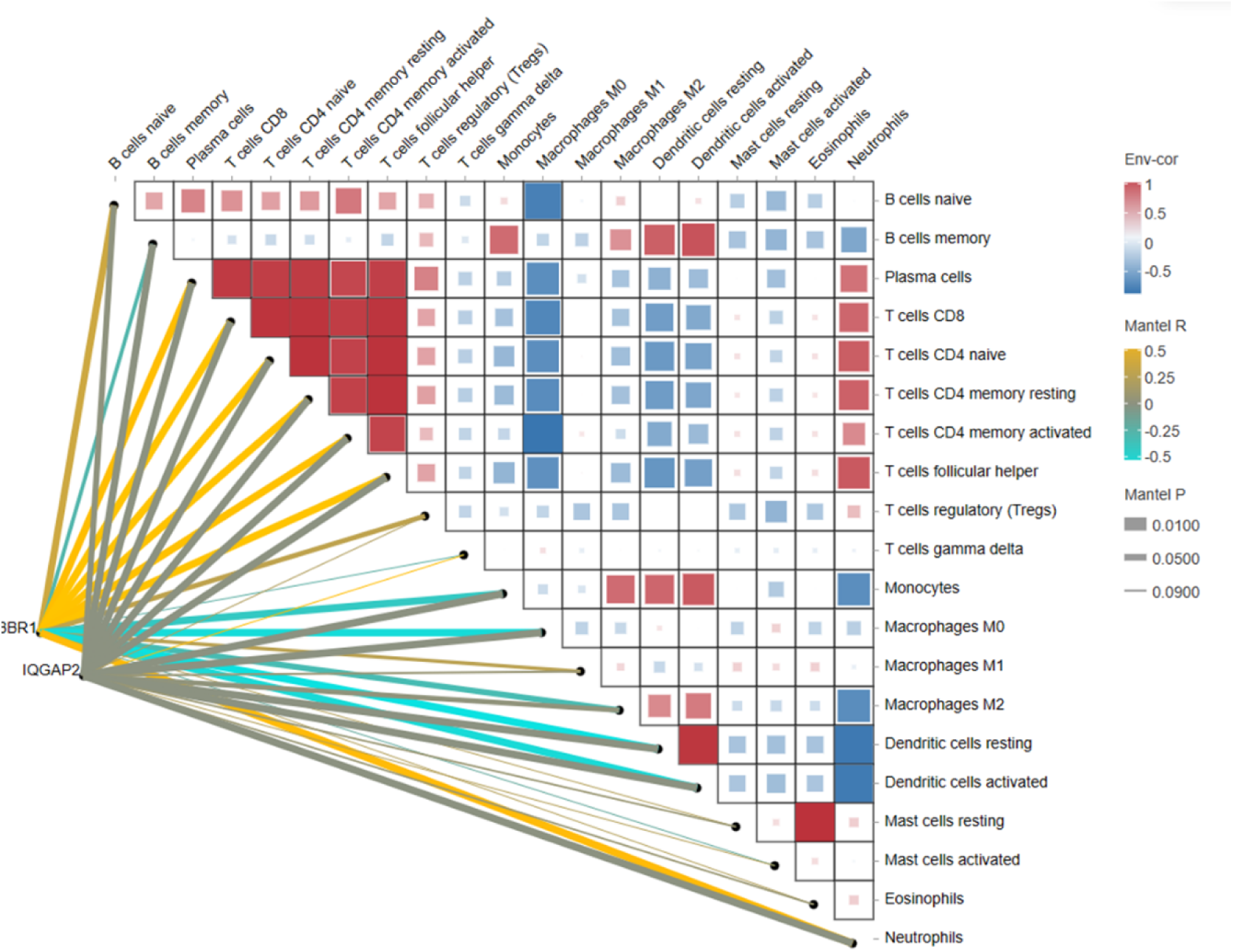
Heat map of the correlation analysis between core genes and immune cells.

### Regulatory miRNA analysis of hub genes

We predicted the target miRNA of hub genes and found that GABBR1 had more potential miRNA than IQGAP2, with a total of 9 miRNAs, while IQGAP2 had only 5 potential miRNAs. Among them, hsa-miR-138-5p and hsa-miR-20a-5p were the co-regulatory target miRNA of the two hub genes (*Figure* 8).

**Figure 8.**
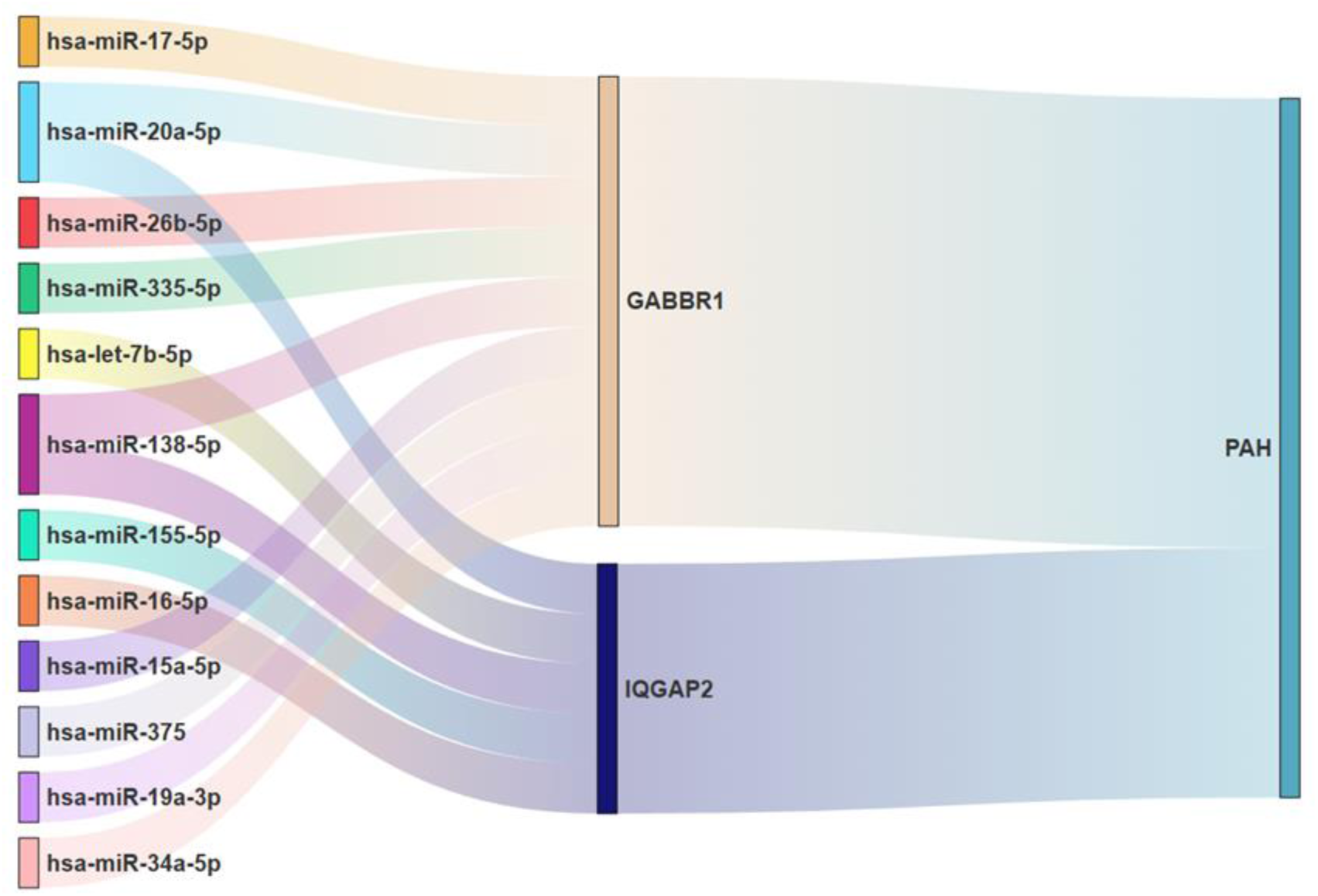
Regulatory miRNAs for hub genes.

## 4. Discussion

The development of PAH is mediated by a combination of genes, and inhibition of the expression of some pivotal genes can effectively intervene in the pathogenesis of PAH (17). The immune infiltration of lung tissue in PAH patients is intrinsically linked to their risk level and prognosis (18). However, the expression of some specific genes can significantly alter the immune cell composition of lung tissue (19). Therefore, it is particularly important to search for pivotal genes in PAH and their relevance to immune infiltration.

### 4.1 Key Findings

In our study, analysis of high-throughput sequencing results from the lungs of PAH patients yielded 132 DEGs, whose functional enrichment revealed that a proportion of DEGs are involved in neuroreceptor activity pathways and are closely associated with immune responses. To further mine the core genes, three machine learning models were constructed. The results revealed that gamma-aminobutyric acid type B receptor subunit 1 (GABBR1) and IQ Motif Containing GTPase Activating Protein 2 (IQGAP2) play key roles in PAH. By validation in peripheral blood and lung tissue, these two pivotal genes could predict the occurrence of PAH. We also analyzed the correlation between the two pivotal genes and immune cells and found that GABBR1 was significantly positively correlated with T-cell populations and negatively correlated with macrophage and dendritic cell populations.

### 4.2 Strengths and limitations

Our study was effective in narrowing the scope of the study by deeply mining the core genes of PAH occurrence through multiple machine learning. The 2 hub genes identified in this study have some ability to predict PAH and may be potential therapeutic targets for PAH. However, there are still some limitations in this study. First, idiopathic PAH can be caused by a variety of factors, such as genetics, residential environment, medications, and strenuous exercise, which were not included in the analysis. Secondly, the subsequent progression of PAH has not been investigated in this study, and the molecular mechanisms underlying the progression of PAH to right heart failure have not been clarified, which has limitations for future clinical application.

### 4.3 Comparison with similar researches

Neuroreceptor activity has long been a hot area of research in PAH. Clinical studies have shown that denervation can effectively improve the morbidity of PAH patients (20). It has been proposed that sympathetic overactivity plays a key role in exacerbating the symptoms of PAH patients (21, 22). In contrast, GABA can effectively inhibit sympathetic nerves and has an important role in maintaining stable sympathetic inhibitory activity (22). This suggests that GABA receptors have a key role in PAH, but the exact mechanism of their regulation is not yet understood.

Pulmonary artery remodeling is an important risk factor for pulmonary arterial hypertension (PAH). The proliferation of arterial endothelium in PAH abnormalities can be attenuated by suppressing the expression of some genes (23). Numerous studies have shown that IQGAP2 is a central regulatory gene of cell proliferation, which promotes cell division by regulating the mTOR pathway (24). However, whether it regulates pulmonary artery remodeling in patients with PAH has not been investigated in depth.

### 4.4 Explanations of findings

Our discovery of the GABBR1 and IQGAP2 genes, which have not been reported in PAH models, could be the focus of subsequent studies. GABBR1 is one of the specific receptors forγ-aminobutyric acid (GABA), which receives GABA signals and thus inhibits neural activity (25). It has been shown that there is a strong link between GABBR1 expression and lung function (26). Sympathetic hyperactivity is an important factor in the pathogenesis of PAH, therefore the expression level of GABBR1 may be related to PAH progression and is a potential therapeutic target for PAH (27). In contrast, IQGAP2 has been shown to have a proliferative effect, promoting angiogenesis and reorganization (28). By down-regulating IQGAP2 it is expected to reduce the state of pulmonary artery remodeling in PAH and thus improve the outcome of PAH patients.

The progression of PAH is closely linked to the immune cell population, particularly the T-cell population, which has a huge potential impact on PAH (29). Among these is the CD4+ subpopulation of T cells (CD4+ T) which functions to resist the inflammatory response of other immune cells, and regulatory T cell deficiency can lead to vascular inflammation with PAH (30). In a lung environment deficient in T cells, lung tissue is more susceptible to vascular endothelial transformation, smooth muscle cell proliferation and outer membrane fibroblast proliferation, which progresses to pulmonary vascular disease (31). In addition to this, studies have shown that regulatory T-cell-deficient rats exhibit accumulation of macrophages a week before and after the onset of PAH (13). In contrast, macrophages secrete Leukotriene B4, a key compound in PAH pulmonary vascular endothelial injury (32). This suggests that a reduced proportion of T cells with the accumulation of macrophages is a feature of the immune infiltration of PAH lung tissue.

We also noted that dendritic cells were recruited in large numbers in the lung tissue of PAH (33). Dendritic cell populations promote the production of pro-inflammatory cytokines such as IL-6, IL-8 and IL-10, and the increase in these inflammatory factors reduces survival in patients with PAH (34, 35). The above findings suggest that the ratio of regulatory T cells, macrophages and dendritic cells is closely related to the progression of PAH, as was also observed in our study. Our study also indicated that GABBR1 expression showed a significant positive correlation with T-cell populations and a significant negative correlation with macrophages and dendritic cells. This suggests that elevating GABBR1 expression may improve the immune infiltration status of lung tissue in PAH.

### 4.5 Implications and actions needed

Traditional PAH biologic factors, such as BMPR2, are highly valuable in predicting the diagnosis and severity of PAH and are widely used in clinical work (36). However, the episodic rate of BMPR2 mutants is low, with only 30% of carriers showing disease manifestations, and some specificity is still lacking in idiopathic PAH (37). Therefore, the exploration and discovery of more specific biological factors is beneficial for the early screening and diagnosis of PAH. In addition, there is an urgent need to develop new therapeutic targets for the treatment of PAH. The new horizons provided by this study will provide some reference for subsequent PAH research.

## 5. Conclusions

Our study identified a total of 2 potential key genes, GABBR1 and IQGAP2, two pivotal genes with good diagnostic significance. They have great potential in regulating neuroreceptor activity and vascular endothelial proliferation in PAH lung tissue. One of them, GABBR1, also has a greater association with the immune infiltration of PAH lung tissue. Our findings may provide a theoretical basis for the analysis of the intrinsic mechanisms of PAH and the development of targeted medicines.

## Acknowledgements

We thank the contributors who previously uploaded and made the dataset publicly available in the GEO database, and all the authors who participated in this study.

## References

1. Ruopp NF, Cockrill BA. Diagnosis and Treatment of Pulmonary Arterial Hypertension: A Review. JAMA 2022;327(14):1379–1391.

2. Simonneau G, Montani D, Celermajer DS, et al. Haemodynamic definitions and updated clinical classification of pulmonary hypertension. Eur Respir J 2019;53(1):1801913.

3. Hoeper M M; Humbert M; Souza R;, et al. A global view of pulmonary hypertension. Lancet Respir. Med. 2016;4:306–322.

4. Frost A, Badesch D, Gibbs JSR, et al. Diagnosis of pulmonary hypertension. Eur Respir J 2019;53(1):1801904.

5. Galiè N, Channick RN, Frantz RP, et al. Risk stratification and medical therapy of pulmonary arterial hypertension. Eur Respir J 2019;53(1):1801889.

6. Suzuki Y J; Nikolaienko S I; Shults N V;, et al. COVID-19 patients may become predisposed to pulmonary arterial hypertension. Med. Hypotheses 2021;147:110483.

7. Shah A J; V orla M; Kalra DK. Molecular Pathways in Pulmonary Arterial Hypertension. Int. J. Mol. Sci. 2022;23:10001.

8. Yildiz, P. Molecular mechanisms of pulmonary hypertension. Clin. Chim. Acta 2009;403:9–16.

9. Bisserier M, Mathiyalagan P, Zhang S, et al. Regulation of the Methylation and Expression Levels of the BMPR2 Gene by SIN3a as a Novel Therapeutic Mechanism in Pulmonary Arterial Hypertension. Circulation 2021;144(1):52–73.

10. Newman J H; Wheeler L; Lane K B;, et al. Mutation in the Gene for Bone Morphogenetic Protein Receptor II as a Cause of Primary Pulmonary Hypertension in a Large Kindred. N. Engl. J. Med 2001;345:319–324.

11. Rabinovitch M, Guignabert C, Humbert M, et al. Inflammation and immunity in the pathogenesis of pulmonary arterial hypertension. Circ Res 2014;115(1):165–175.

12. Huertas A, Tu L, Gambaryan N, et al. Leptin and regulatory t-lymphocytes in idiopathic pulmonary arterial hypertension. Eur Respir J 2012;40:895–904.

13. Tamosiuniene R, Tian W, Dhillon G, et al. Regulatory t cells limit vascular endothelial injury and prevent pulmonary hypertension. Circulation research 2011;109:867–879.

14. Steiner MK, Syrkina OL, Kolliputi N, et al. Interleukin-6 overexpression induces pulmonary hypertension. Circulation research 2009;104:236–244.

15. Barrett T, Wilhite SE, Ledoux P, et al. NCBI GEO: archive for functional genomics data sets--update. Nucleic Acids Res 2013;41(Database issue):D991–995.

16. Newman AM, Liu CL, Green MR, et al. Robust enumeration of cell subsets from tissue expression profiles. Nat Methods 2015;12(5):453–457.

17. Li D, Shao N Y, Moonen J R, et al. ALDH1A3 Coordinates Metabolism With Gene Regulation in Pulmonary Arterial Hypertension. Circulation 2021;143(21):2074–2090.

18. Wang R R, Yuan T Y, Wang J M, et al. Immunity and inflammation in pulmonary arterial hypertension: From pathophysiology mechanisms to treatment perspective. Pharmacol Res 2022;180:106238.

19. Yaku A, Inagaki T, Asano R, et al. Regnase-1 Prevents Pulmonary Arterial Hypertension Through mRNA Degradation of Interleukin-6 and Platelet-Derived Growth Factor in Alveolar Macrophages. Circulation 2022;146(13):1006–1022.

20. Zhang H, Wei Y, Zhang C, et al. Pulmonary Artery Denervation for Pulmonary Arterial Hypertension: A Sham-Controlled Randomized PADN-CFDA Trial. JACC Cardiovasc Interv 2022;15(23):2412–2423.

21. Simpson L L, Meah V L, Steele A, et al. Evidence for a physiological role of pulmonary arterial baroreceptors in sympathetic neural activation in healthy humans. J Physiol 2020;598(5):955–965.

22. Yu Q, Guo Q, Jin S, et al. Melatonin suppresses sympathetic vasomotor tone through enhancing GABAA receptor activity in the hypothalamus. Front Physiol 2023;14:1166246.

23. Zhang L, Wang Y, Wu G, et al. Blockade of JAK2 protects mice against hypoxia-induced pulmonary arterial hypertension by repressing pulmonary arterial smooth muscle cell proliferation. Cell Prolif 2020;53(2):e12742.

24. Chen T, Fan X, Li G, et al. Multi-omics data integration reveals the molecular network of dysregulation IQGAP2-mTOR promotes cell proliferation. Hum Cell 2023;36(4):1429–1440.

25. Enoch M A, Hodgkinson C A, Shen P H, et al. GABBR1 and SLC6A1, Two Genes Involved in Modulation of GABA Synaptic Transmission, Influence Risk for Alcoholism: Results from Three Ethnically Diverse Populations. Alcohol Clin Exp Res 2016;40(1):93–101.

26. George L, Mitra A, Thimraj T A, et al. Transcriptomic analysis comparing mouse strains with extreme total lung capacities identifies novel candidate genes for pulmonary function. Respir Res 2017;18(1):152.

27. Zhang H, Chen S L. Pulmonary Artery Denervation: Update on Clinical Studies. Curr Cardiol Rep 2019;21(10):124.

28. Jahejo A R, Rajput N, Kashif J, et al. Recombinant glutathione-S-transferase A3 protein regulates the angiogenesis-related genes of erythrocytes in thiram induced tibial lesions. Res Vet Sci 2020;131:244–253.

29. Meng X, Yang J, Dong M, et al. Regulatory T cells in cardiovascular diseases. Nat Rev Cardiol 2016;13(3):167–179.

30. Qiu H, He Y, Ouyang F, et al. The Role of Regulatory T Cells in Pulmonary Arterial Hypertension. J Am Heart Assoc 2019;8(23):e014201.

31. Qian J, Tian W, Jiang X, et al. Leukotriene B4 Activates Pulmonary Artery Adventitial Fibroblasts in Pulmonary Hypertension. Hypertension 2015;66(6):1227–1239.

32. Tian W, Jiang X, Tamosiuniene R, et al. Blocking macrophage leukotriene b4 prevents endothelial injury and reverses pulmonary hypertension. Sci Transl Med 2013;5(200):200ra117.

33. Perros F, Dorfmuller P, Souza R, et al. Dendritic cell recruitment in lesions of human and experimental pulmonary hypertension. Eur. Respir. J 2007;29:462–468.

34. George P M, Badiger R, Shao D, et al. Viral Toll Like Receptor activation of pulmonary vascular smooth muscle cells results in endothelin-1 generation; relevance to pathogenesis of pulmonary arterial hypertension. Biochem. Biophys. Res. Commun 2012;426:486–491.

35. Soon E, Holmes A M, Treacy C M, et al. Elevated levels of inflammatory cytokines predict survival in idiopathic and familial pulmonary arterial hypertension. Circulation 2010;122:920–927.

36. Tatius B, Wasityastuti W, Astarini FD, et al. Significance of BMPR2 mutations in pulmonary arterial hypertension. Respir Investig 2021;59(4):397–407.

37. Song J, Hinderhofer K, Kaufmann L T, et al. BMPR2 Promoter Variants Effect Gene Expression in Pulmonary Arterial Hypertension Patients. Genes (Basel) 2020;11(10):1168.

